# Transfer Learning Models for Bacterial Strain Dissemination Biomarkers using Weighted Non-Parallel Proximal Support Vector Machines

**DOI:** 10.1101/2024.10.11.617744

**Authors:** Ugochukwu O. Ugwu, Richard A. Slayden, Michael Kirby

## Abstract

This paper develops optimization and Machine Learning (ML) algorithms to analyze gene expression datasets from the lungs and spleen of mice, infected intranasally, with two bacterial strains, *Francisella tularensis* - Schu4 and Live Vaccine Strain (LVS). We propose and utilize Weighted *𝓁*_1_-norm Generalized Eigenvalue-type Problems (*𝓁*_1_-WGEPs) to determine a small set of host biomarkers that report Schu4 and LVS infection of the lungs and dissemination to the spleen. The optimal solutions of *𝓁*_1_-WGEPs determine the direction onto which the datasets are projected for dimensionality reduction, with the projection scores computed and ranked for gene selection. The top *k*-ranked projection scores correspond to the top *k* most informative biomarker features. The top *k* features selected from the lungs data are employed to train ML models, with uninfected controls and Schu4 or LVS samples as classes. The trained models are validated on the spleen data to incorporate transfer learning. Baseline ML algorithms such as ANN, XGBoost, AdaBoost, AdaGrad, KNN, SVM, Naive Bayes, Random Forest, Logistic Regression, and Decision Tree are compared with our Weighted *𝓁*_1_-norm Non-Parallel Proximal Support Vector Machine (*𝓁*_1_-WNPSVM) that is based on two non-parallel separating hyperplanes. We report average balanced accuracy scores of the methods over multiple folds. Gene ontology is performed on the most significant genes in both tissues to reveal biomarkers of disease and examine for relevant metabolic pathways for host-directed therapeutics development and treatment performance.

**Author Summary:** Integrating genomic datasets from homogeneous or heterogeneous sources is an area that is currently underexplored. This work develops new methodologies to integrate transcriptomic datasets from the lungs and spleen tissues infected by Francisella tularensis — Schu4 and Live Vaccine Strain (LVS). Our objective is to identify biologically relevant gene features indicative of respiratory infection, disease severity, and bacterial dissemination to the spleen, then utilize the selected features to predict disease status using our Weighted *𝓁*_1_-norm Non-Parallel Support Vector Machines (*𝓁*_1_-WNPSVM), which is trained on the lungs data and validated on the spleen data, introducing a form of transfer learning. The *𝓁*_1_-WNPSVM outperforms traditional ML techniques, achieving a 97% balanced accuracy. It also generalizes to models of similar formulations, incorporating dimensionality reduction and gene selection into the NPSVM-type framework. Currently, a direct application of existing NPSVM-type methods to analyze gene expression datasets, where the number of genes significantly exceeds the number of samples, is computationally impractical due to their large memory requirements. This work addresses this challenge. We discovered sets of 253 genes exclusively expressed in the lungs and spleen tissues. Gene ontology is performed to reveal underlying metabolic pathways. Our analysis shows that the immune system pathway is activated in both lungs and spleen.

## 1 Introduction

This paper develops novel optimization and Machine Learning (ML) algorithms involving transfer learning. The importance of applying optimization and ML algorithms with transfer learning to biological problems lies in improving model performance, scaling across applications, and complex problem-solving. Utilization of this approach will facilitate our goal to identify host biomarkers of bacterial respiratory infections, elucidate dissemination patterns, and uncover underlying metabolic pathways of the host-pathogen interactions. This development is achieved by analyzing gene expression datasets, from the lungs and spleen tissues of genetically identical mice, infected intranasally, with two bacterial strains, *Francisella tularensis* - Schu4 and Live Vaccine Strain (LVS). We anticipate that our approach will advance biomarker and biological pathway discovery and aid in the development of therapeutics with efficacy against difficult-to-treat medically important pathogens and cancer.

*Francisella tularensis* is the causative agent of tularemia, also known as *rabbit fever*. According to the U.S. Centers for Disease Control and Prevention (CDC), tularemia is a rural disease that has been reported in all U.S. states except Hawaii^1^. *Francisella tularensis* is highly virulent with bioterrorism potential even at a very low dose. *Francisella tularensis* Schu4 strain is a highly virulent Type A strain that causes severe disseminated disease in humans and animals. *Francisella tularensis* LVS strain is a less virulent attenuated Type B strain that causes a milder form of disease. These strains are routinely used in drug and vaccine research studies because they are genetically stable and have well-defined coding capacity and virulence factors, and importantly, afford the opportunity to study mechanisms involved in the development of severe and mild diseases as well as dissemination.

### 1.1 Our Contributions

Schu4 induces a robust immune response due to its high virulence, making it useful for studying host-pathogen interactions and immune evasion mechanisms while LVS elicits a less intense immune response resulting in different levels of virulence. Accordingly, we have chosen to utilize host transcriptional data from infected lungs and spleen to define a set of biomarkers from biologically relevant gene features indicative of respiratory infection, disease severity and bacterial dissemination to the spleen.

Here and throughout this paper, we will use the matrices *𝒢*_1_∈ ℝ^*m×n*^ and *𝒢*_2_ ∈ ℝ^*m×n*^ to represent gene expression datasets from the lungs and spleen tissues, respectively, where each matrix consists of *m* genes and *n* samples. The columns of *𝒢*_1_ and *𝒢*_2_ will include controls, Schu4 samples, and LVS samples, in that order. The class matrices, 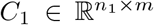 and 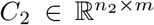, in our context, represent all the training data points from *𝒢*_1_ or *𝒢*_2_ that correspond to Class 1 (uninfected controls) and Class –1 (infected Schu4 or LVS samples), respectively, with *n > n*_1_ + *n*_2_. The variables *n*_1_ and *n*_2_ represent the number of samples in each class.

Also, as with most transcriptional data, *𝒢*_1_∈ ℝ ^*m×n*^ and *𝒢*_2_ ∈ ℝ ^*m×n*^ are high-dimensional and have significantly more genes than samples, i.e., *m* ≫ *n*. Thus, necessitating the need for gene selection, a process that reduces the number of genes by removing redundant and irrelevant genes. Notably, the gene selection methods can be categorized into supervised approach utilizing label information, unsupervised approach involving the use of unlabeled data to identify patterns or structures within the dataset, and semi-supervised or semi-unsupervised approach relying on both labeled and unlabeled data. For more discussions on gene selection techniques, see, e.g., [35] and references therein. We will focus on identifying the most informative gene expression profiles that potentially reflect host immune responses to bacterial dissemination across tissues, then use the selected group of genes to carry out binary classification tasks.

Our approach to gene selection and classification involves modifying the *𝓁*_1_-norm generalized eigenvalue-type *Non-Parallel Proximal Support Vector Machine* [4, *𝓁*_1_-NPSVM] to make it applicable to transcriptional data analysis. Currently, a direct application of existing *𝓁*_1_-NPSVM architecture [2, 3] to gene expression datasets, is computationally impractical due to their large memory requirements. Specifically, the solution methods [2, 3] involve large-scale matrix-matrix computations, with 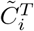 and with 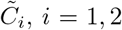, *i* = 1, 2, at each iteration, thus resulting in matrices of size (*m* + 1) × (*m* + 1). This makes solving (1.1) infeasible for simultaneous gene selection and classification tasks. Moreover, the computational cost at each iteration is doubled especially since problems (1.1) are solved concurrently.

The *𝓁*_1_-NPSVM formulation was first proposed in [4], and it is given by the following large-scale minimization problems

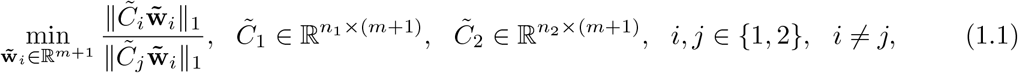

where 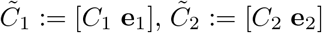, and 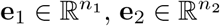 are vectors of ones. The quantity ∥ · ∥_1_ denotes the *𝓁*_1_-norm of a vector **y** given by

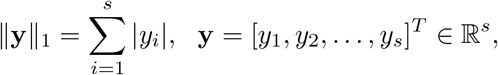

where |*y*| denotes the absolute value of *y* ∈ R. Additionally, the solutions of (1.1), 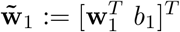 and 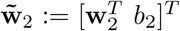, determine two non-parallel separating hyperplanes,

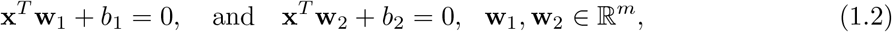

where *b*_1_, *b*_2_ ∈ ℝ are the *bias* terms and **x** ∈ ℝ^*m*^ is a new sample point to be classified to either Class 1 (*C*_1_) or Class -1 (*C*_2_) according to (2.24). The superscript ^*T*^ denotes transposition.

The contributions of this paper can be summarized in fourfold:

1. We propose the use of weighted *𝓁*_2_-norm for the regularization of (1.1). This leads to the minimization problems

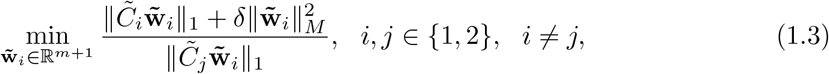

where

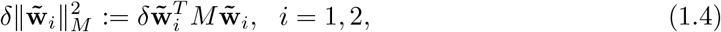

and *M* ∈ ℝ^(*m*+1)*×*(*m*+1)^ is a symmetric positive definite (SPD) matrix or a diagonal matrix different from the identity matrix *I*. The quantity 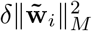, *i* = 1, 2, are referred to as the *regularization terms*, and for a fixed *i* and *j* with *i* ≠ *j, δ >* 0 is a regularization parameter that balances the influence of 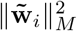 on the solutions of (1.3). A common approach to determine *δ* is by cross-validation or grid search. Our approach of regularizing (1.1) is consistent with the Grassmannian regularization which specifies *M* in relation to the *Graph Laplacian Matrix*, see, e.g., [17, 18]. Also, by following [2, 3], we can express (1.1) as constrained minimization problems

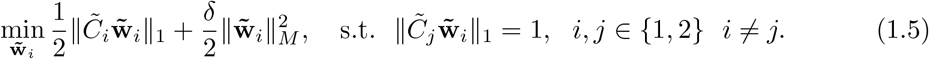

We will refer to (1.5) as the Weighted *𝓁*_1_-NPSVM (*𝓁*_1_-WNPSVM). When *M* = *I*, the resulting problems, referred to as the *𝓁*_1_-GEPSVM [3], are special cases of the *𝓁*_*q*_-NPSVM, *q >* 0, which is described in [2]. The *𝓁*_*q*_-NPSVM regularizes (1.1) by adding 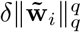 to the numerator. We will show in Section 2 that 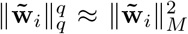 for *q >* 0, where *M* is a diagonal matrix that changes at each iteration and also depends on *q* and 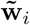, *i* = 1, 2. Other related but different regularization techniques are presented in, e.g., [15, 19, 20, 21]. We will show in Section 2 that our approach generalizes the regularization methods described in [2, 3].
2. We introduce new methodologies that integrate dimensionality reduction and feature selection into the *𝓁*_1_-WNPSVM framework. This is to enable practical application to gene expression datasets, particularly in situations where the number of genes significantly exceeds the number of samples (*m* ≫ *n*). The dimensionality reduction of *𝒢*_1_ and *𝒢*_2_ is accomplished by solving constrained Weighted *𝓁*_1_-norm Generalized Eigenvalue-type Problems (*𝓁*_1_-WGEPs)

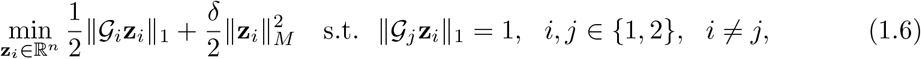

where *M* ∈ ℝ^*n×n*^ is a SPD matrix that may depend on **z**_*i*_, *i* = 1, 2. After solving (1.6), *𝒢*_1_ and *𝒢*_2_ are projected onto one-dimensional subspaces spanned by the solutions, **z**_1,*δ*_ and **z**_2,*δ*_, to identify *k*≪ *m* most informative genes. This reduces *𝒢*_1_ and *𝒢*_2_ to small size matrices of dimensions *k* × *n*, that are suitable for use in (1.5) for binary classification. The minimization problems (1.6) can be thought of as *unsupervised multiclass gene selection problems* since each *𝒢*_*i*_, *i* = 1, 2, consist of controls, Schu4 samples and LVS samples as classes, and the most informative genes across the classes are selected synchronously.
3. The projection directions are determined by the solutions of (1.6). The best direction for each dataset is determined by computing the maximum projection scores of *𝒢*_1_ and *𝒢*_2_ with respect to **z**_1,*δ*_ and **z**_2,*δ*_, i.e.,

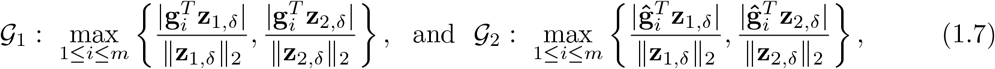

where **g**_*i*_, **ĝ**_*i*_ ∈ ℝ^*n*^, *i* = 1, 2, …, *m*, are the gene expression vectors in *𝒢*_1_ and *𝒢*_2_, respectively. Note that a direction that gives the maximum score for each dataset is used for gene selection. We will select *k*≪ *m* most significant genes from *𝒢*_1_ and *𝒢*_2_ by computing their projection scores with a chosen projection direction, where the top *k* projection scores correspond to the top *k* most informative genes. Details of dimensionality reduction and gene selection are provided by Algorithms 1 and 2 in Section 2.
4. We discovered sets of 253 genes that are uniquely expressed in the lungs and spleen tissues. Notably, the gene sets from the lungs and spleen do not overlap. Among these genes, only 221 and 190 gene identifiers from the lungs and spleen, respectively, are available for transcriptomic pathway analysis in Reactome. Our functional enrichment analyses identified 159 of the 221 identifiers and 137 of the 190 identifiers that are linked to 776 and 1023 pathways, respectively. The top 10 most significant pathways activated in the lungs are associated with essential functions in developmental biology, regulation of immune system cells development, signal transduction, disease (cancer), and metabolism pathways, while the leading pathways in the spleen comprise of signal transduction, immune system, disease (tuberculosis), developmental biology, and transport of molecules pathways. A more elaborate discussion focusing on pathways discovery is presented in Section 3.

In the remainder of this section, we will provide some details on related methods.

### 1.2 Related Methods

The idea to use two non-parallel separating hyperplanes for binary classification tasks was first proposed in a seminal work by Mangasarian and Wild [5]. It requires that each hyperplane be closest to the samples of one class and farthest from the samples of the other class. Therefore, the assignment of **x** to *C*_1_ or *C*_2_ depends on the hyperplane it is most proximal to.

Several different formulations of (1.1) have been proposed and applied in the literature, with binary classification applications to functional data [6], crossed data [5], satellite ship image data [7], ellipsoidal uncertainty data [8], gene expression data [9] including genomic and proteomic problems [10], biomedical data [11], lungs cancer cells [12], as well as applications to multiclass [5] and multiview learning [13] problems. It is noteworthy that the (regularized) *𝓁*_1_-NPSVM has been applied to binary classification tasks in [2, 3, 4], however, this paper is the first to apply (1.1) in the context of microarray or transcriptomic pattern analysis. Our approach differ significantly from the models described in [9].

The use of *𝓁*_1_-norm in (1.1) instead of squared *𝓁*_2_-norm is to reduce the sensitivity of the solutions to noise and outliers. The minimization problems (1.1) with the squared *𝓁*_2_-norm are discussed in [5]. They are commonly referred to as the *Generalized Eigenvalue Proximal Support Vector Machine* (GEPSVM). The GEPSVM results in the *Rayleigh Quotients*,

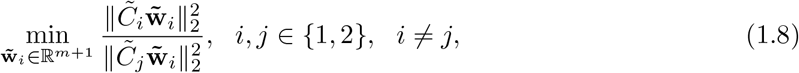

where for any matrix *A* and a vector **x** of compatible sizes, 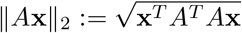. The solutions of (1.8) are readily determined by solving the generalized eigenvalue problems

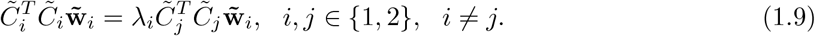

The eigenvectors corresponding to the smallest eigenvalues of 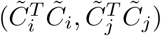, for *i, j* = 1, 2, *≠ j*, are the optimal solutions of (1.8). Also, the objective functions in (1.8) share the same eigenvectors but have reciprocal eigenvalues 1*/λ*_*i*_. Therefore, the eigenvector corresponding to the largest eigenvalue of (1.9) when *i* = 1 and *j* = 2 is also an eigenvector corresponding to the smallest eigenvalue of (1.9) when *i* = 2 and *j* = 1.

Additionally, the Rayleigh quotients in (1.8) are bounded and ranges over an interval determined by their minimum and maximum eigenvalues when the denominator is positive definite [14]. However, the defining matrices in (1.8) are rank-deficient, as they have a maximum rank of *m* despite being of order *m* + 1. This makes the minimization problems (1.8) prone to becoming singular, with *N* (*C*_*i*_) ∩ *N* (*C*_*j*_) ≠ **0**, where *N* (*A*) denotes the null space of the matrix *A* and **0** is the zero vector [15]. Consequently, even when (1.8) is defined by using positive semi-definite matrices, it often fails to produce meaningful approximations of the generalized eigenvectors [2]. As a remedy to the difficulty in solving (1.8), [5] proposed a solution approach that replaces (1.8) with nearby problems that are more stable to solve. Specifically, they consider regularized minimization problems

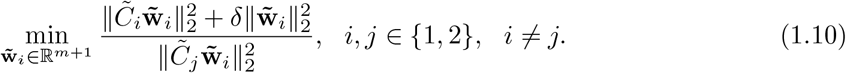

The replacement of (1.8) by (1.10) is known as Tikhonov regularization [16] in *standard form*. It should be noted that although the squared *𝓁*_2_-norm regularization terms in (1.10) can improve the stability and classification accuracy of the GEPSVM, it may also amplify the influence of outliers in the samples and cause decreased accuracy. In comparison, the *𝓁*_1_-NPSVM has been found to be more robust to outliers than the GEPSVM, as discussed in [2, 3]. Moreover, the GEPSVM has been demonstrated in [5] to yield better accuracy than the Support Vector Machine [30, SVM] when applied to solve classical ‘XOR’ and ‘Cross Planes’ problems.

### 1.3 Organization of this work

This paper is organized as follows. Section 2 discusses the solution methods for the minimization problems (1.5) and (1.6), and describes how (1.6) can be applied to achieve dimensionality reduction and gene selection. The data integration process involves fusing uninfected (controls), Schu4 samples, and LVS samples from the lungs and spleen tissues. Once a small set of genes that indicate bacterial dissemination of Schu4 and LVS strains has been identified for both tissues, transfer learning is carried out by using (1.5). Information from the lungs is used to train transfer learning models while the trained models are applied to the spleen data to compute balanced classification accuracy scores. Numerical experiments of Section 3 discuss the performance of various ML models, such as Artificial Neural Networks (ANN) [22], Adaptive Boosting (AdaBoost) [23], Gradient Boosting (GradBoost) [24], Extreme Gradient Boosting (XGBoost) [25], K-Nearest Neighbors (KNN) [26], Logistic Regression [27], Random Forest [28], Naive Bayes [29], Support vector Machine (SVM), and Decision Tree [31]. These models are trained with uninfected controls and Schu4 or LVS samples as classes and are compared with our proposed *𝓁*_1_-WNPSVM model. In Section 4, transcriptomic pattern analyses of a select group of genes with pathway analysis are presented while Section 5 provides concluding remarks.

## 2 The Solution Methods for (1.5) and (1.6)

This section describes the solution methods for (1.5) and (1.6). Since the problems are analogous, we will instead, without loss of generality, consider the minimization problem

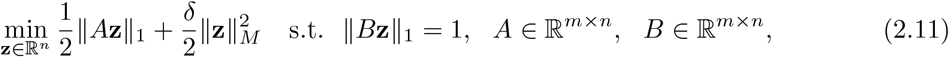

where the vectors **a**_*i*_ ∈ ℝ ^*n*^, and **b**_*i*_ ∈ ℝ ^*n*^, *i* = 1, 2, …, *m*, denote the rows of *A* and *B*, respectively. Here and below, we will use boldface letters, e.g., **a**, to denote vectors and capital letters, e.g., *A*, to denote matrices.

By following [2, 3], we will approximate ∥ · ∥ _1_ by a weighted *𝓁*_2_-norm, then reformulate (2.11) into a compact and stable quadratic minimization problem. Hence, for *ϵ >* 0,

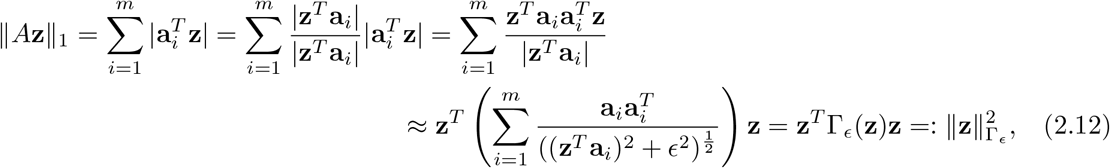

where

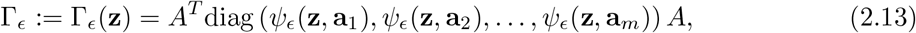

and 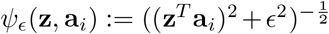. The matrix Γ is symmetric, and the first term in line 2 of (2.12) accounts for the possibility of | **z**^*T*^ **a**_*i*_ |≈ 0, by adding a small parameter *ϵ* to the denominator. This is different from [2, 3] which assumes that | **z**^*T*^ **a**_*i*_| ≠ 0 and expresses ∥*A***z** ∥_1_ as in line 1 of (2.12). Also,

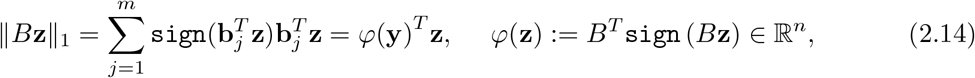

where sign(**x**) is an element-wise sign of a vector **x**. Substitution of (2.12) and (2.14) into (2.11) gives

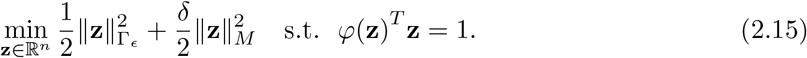

Equation (2.15) leads to the constrained quadratic programming problem

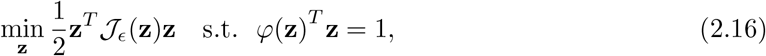

where *𝒯* _*ϵ*_(**z**) = Γ_*ϵ*_(**z**) + *δM*.

We will employ an *iterative optimization strategy* described in [2, 3] to compute the solution of (2.16). The basic idea is as follows. Let **z**^(*k*)^, *k* ≥ 1, be the currently available approximation of the solution

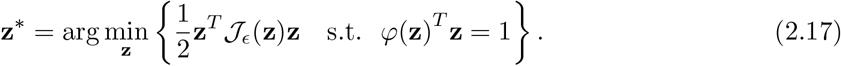

Then

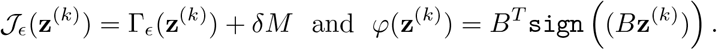

To derive an update for **z**^(*k*+1)^, denote the Langrangian of (2.16) as

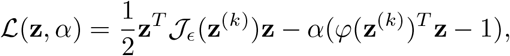

where *α* is the Langrangian multiplier. Then

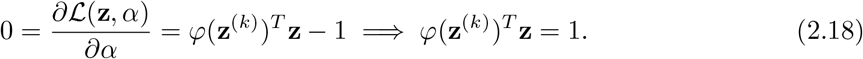

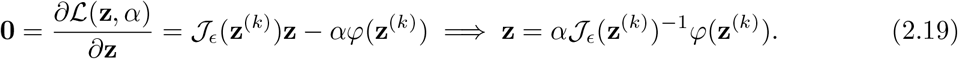

Substitute (2.19) into (2.18) to determine *α*, then use the result in (2.19) to obtain the desired update for **z**^(*k*+1)^ as

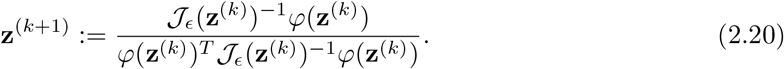

Thus, at each iteration, a new approximation **z**^(*k*+1)^ of **z**^*\**^ is computed by using the current *𝒯* _*ϵ*_(**z**^(*k*)^) and *φ*(**z**^(*k*)^). This process continues repeatedly until certain convergence conditions are met.

The solution method so described is summarized by Algorithm 1. This algorithm solves (2.11) and the minimization problem

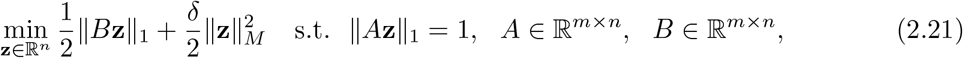

simultaneously, by reversing the roles of *A* and *B* in step 14. The influence of the parameters, *ϵ* and *δ*, will be examined in Section 3. The convergence results for Algorithm 1 are readily shown by following [2, 3].

In the sequel, the following proposition demonstrates that our approach of regularizing (1.1) with a weighted *𝓁*_2_-norm generalizes the regularization methods described in [2, 3].

**Proposition 2.1**. Consider the minimization problem [2]

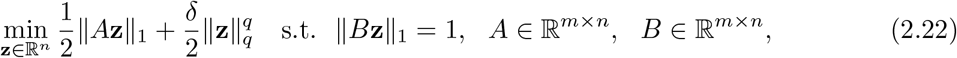

analogous to (2.11) and (2.21). Then (2.22) can be expressed in the form (2.16).

*Proof:* Let 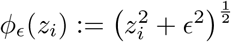 for *ϵ >* 0. Then

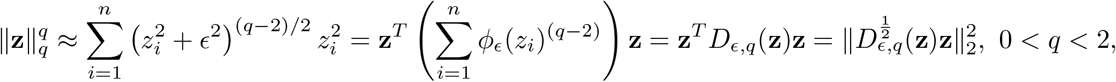

where **z** := [*z*_1_, *z*_2_, …, *z*_*n*_]^*T*^, and

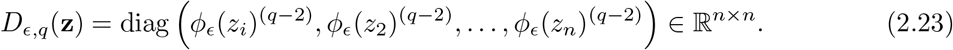

Setting *M* := *D*_*ϵ,q*_(**z**) completes the proof. □

In the computed examples of Section 3, we will compare the performance of *𝓁*_1_-WGEPs for *M* := *D*_*ϵ*,1_(**z**) and *M* := *D*_*ϵ*,1_(**z**)^2^ to illustrate that the latter matrix captures more information than the former in terms of the overall projection scores.

### Algorithm 1: The Solution Method for the Approximate Solution of (2.11) and (2.21)

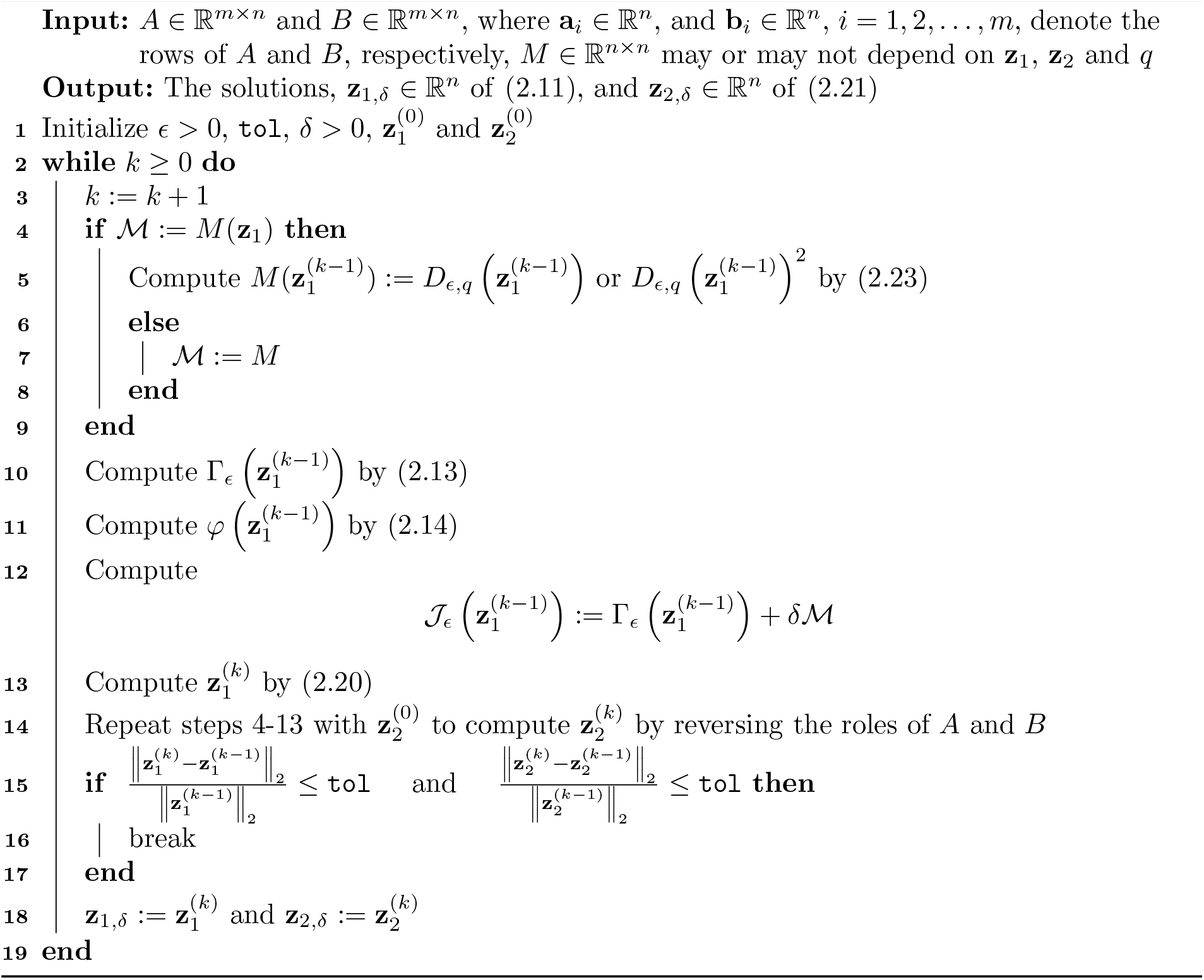

### Algorithm 2: The *𝓁*_1_-WGEPs for Dimensionality Reduction and Gene Selection

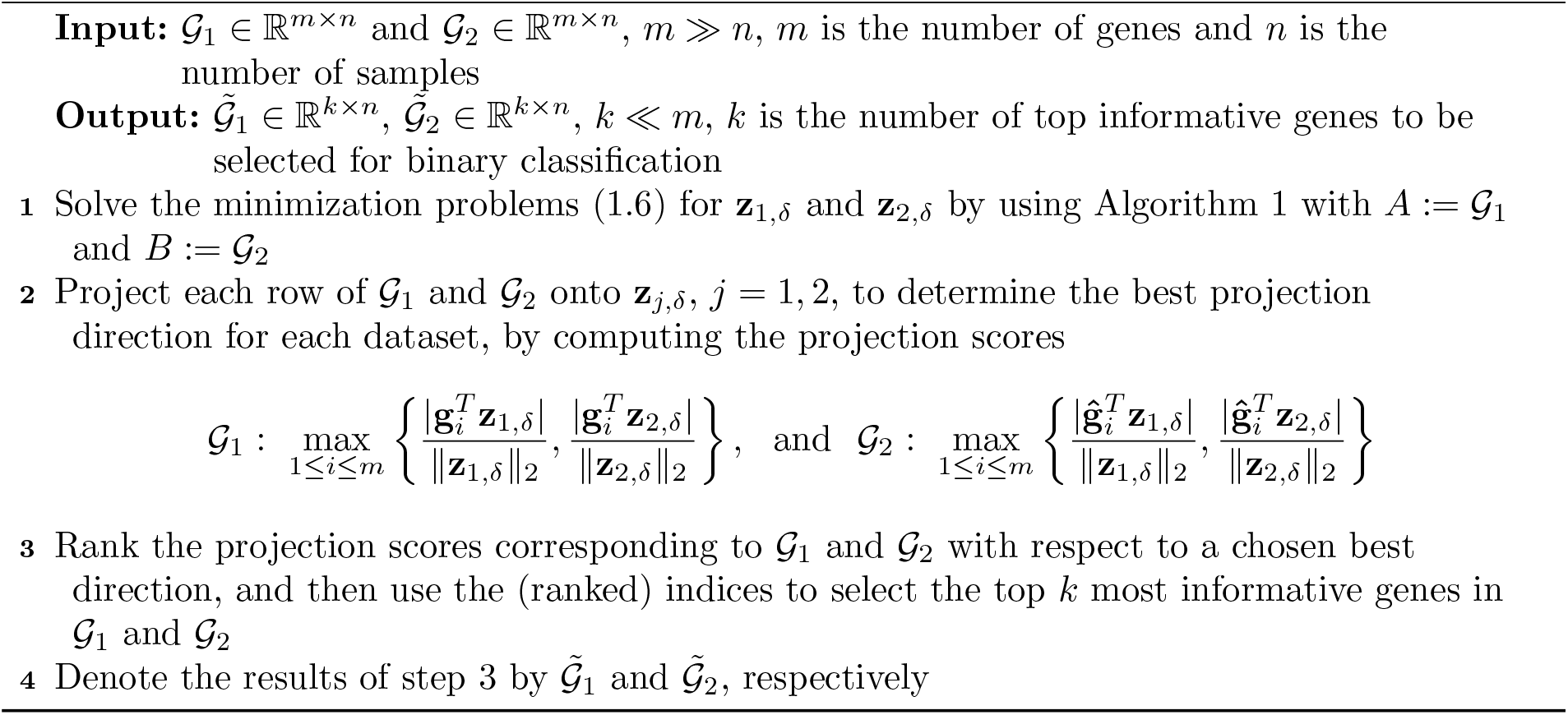

Algorithm 2 describes the *𝓁*_1_-WGEPs for dimensionality reduction and gene selection. By applying Algorithm 2 to *𝒢* _1_ ∈ ℝ ^*m×n*^ and *𝒢* _2_ ∈ ℝ ^*m×n*^, the matrices are reduced to small size matrices, 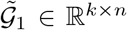, and 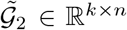, respectively. The rows of 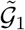 and 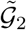 contain the top *k* ≪ *m* most informative genes selected for the binary classification of controls and Schu4 or LVS samples. In other words, to solve (1.5) by Algorithm 3, we use 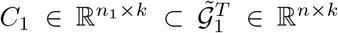, and 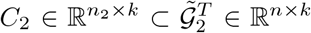, where *n*_1_ + *n*_2_ *< n*. Note that *n*_1_ and *n*_2_ are the number of controls and Schu4 or LVS samples in *𝒢* _1_ or *𝒢* _2_, respectively.

Algorithm 3 describes the solution methods for the proposed *𝓁*_1_-WNPSVM formulations. The outputs of Algorithm 3, **w**_1_, **w**_2_, *b*_1_, and *b*_2_, determine two non-parallel separating hyperplanes (1.2). A new sample point **x** is classified to either Class 1 (*C*_1_) or Class -1 (*C*_2_) by using

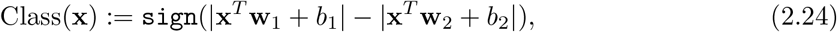

where sign(*x*) is the sign function of *x ∈* ℝ. This is equivalent to

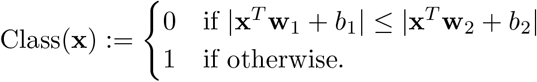

In the following section, we discuss the binary classification of uninfected controls and Schu4 or LVS samples, and compare *𝓁*_1_-WNPSVM method with several baseline algorithms.

### Algorithm 3: The *𝓁*_1_-WNPSVM (1.5) for Binary Classification

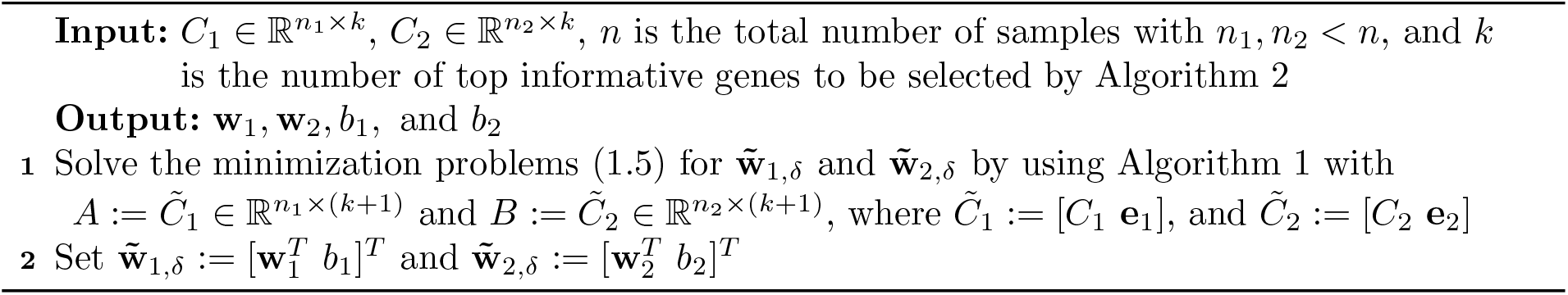

## 3 Numerical Experiments

This section discusses the performance of several baseline ML classifiers, such as Random Forest, Logistic Regression, SVM, Decision Tree, Naive Bayes, KNN, AdaBoost, GradBoost, XGBoost, and ANN, on binary classification problems. These methods will be compared with the *𝓁*_1_-WNPSVM in Jupyter Notebook, version 6.3.0. All computations are carried out on a Dell computer running Windows 11 with 11th Gen Intel(R) Core(TM) i7-1165G7 @ 2.80GHz and 16 GB RAM.

### 3.1 Preprocessing of the Lungs and Spleen DataSets

The lungs and spleen datasets contain 6 controls, 24 Schu4 samples, and 24 LVS samples in that order, resulting in *n* = 54 samples. The samples are taken at four time points, 12, 24, 48, and 120 hours, with a total of *m* = 37, 632 genes measured at each time point under the same experimental conditions for both tissues [1].

The lungs and spleen datasets are imbalanced, with only 6 uninfected controls each. Therefore, to preprocess and integrate both datasets as well as address the imbalance, the *Synthetic Minority Oversampling Technique* [32, SMOTE] is applied to the available 6 controls in other to generate additional 18 synthetic controls. The SMOTE algorithm^2^ applies linear interpolation to synthesize a new control between two control points.

The preprocessing stage is carried out in MATLAB (R2021a). Genes with standard deviations less than 0.9 are thrown out from the lungs and spleen datasets, leaving a total of 19, 735 genes in each dataset. Refer to [1] for a detailed description of the datasets, including normalization.

### 3.2 Integrating the Lungs and Spleen Datasets for Dimensionality Reduction and Gene Selection

The *𝓁*_1_-WGEPs (1.6) are used to integrate the preprocessed lungs and spleen datasets. The goal of data integration is to identify a small group of informative genes that significantly contribute to the process of bacterial dissemination of Schu4 and LVS across tissues, and more so, dimensionally reduce the sizes of the lungs (*𝒢*_1_) and spleen (*𝒢*_2_) datasets.

The solutions of (1.6) are determined by Algorithm 1 with *M* := *D*_*ϵ*,1_(**z**)^2^ and tol = 10^*−*4^, while step 2 of Algorithm 2 or (1.7) determines the best direction to project each dataset for gene selection. The projection directions, **z**_1,*δ*_ and **z**_2,*δ*_, are found to be best for the spleen and lungs datasets, respectively.

Figure 1 displays the projection scores corresponding to *𝒢* _1_ and *𝒢*_2_ for different values of the parameters, *δ* and *ϵ*. It is evident from Figure 1 that *δ* = 1000 and *ϵ* = 10^*−*3^ yield the highest projection scores for both datasets, indicating that the corresponding *𝓁*_1_-WGEPs capture more information than with other *δ* and *ϵ* values shown in Figure 1.

**Figure 1:**
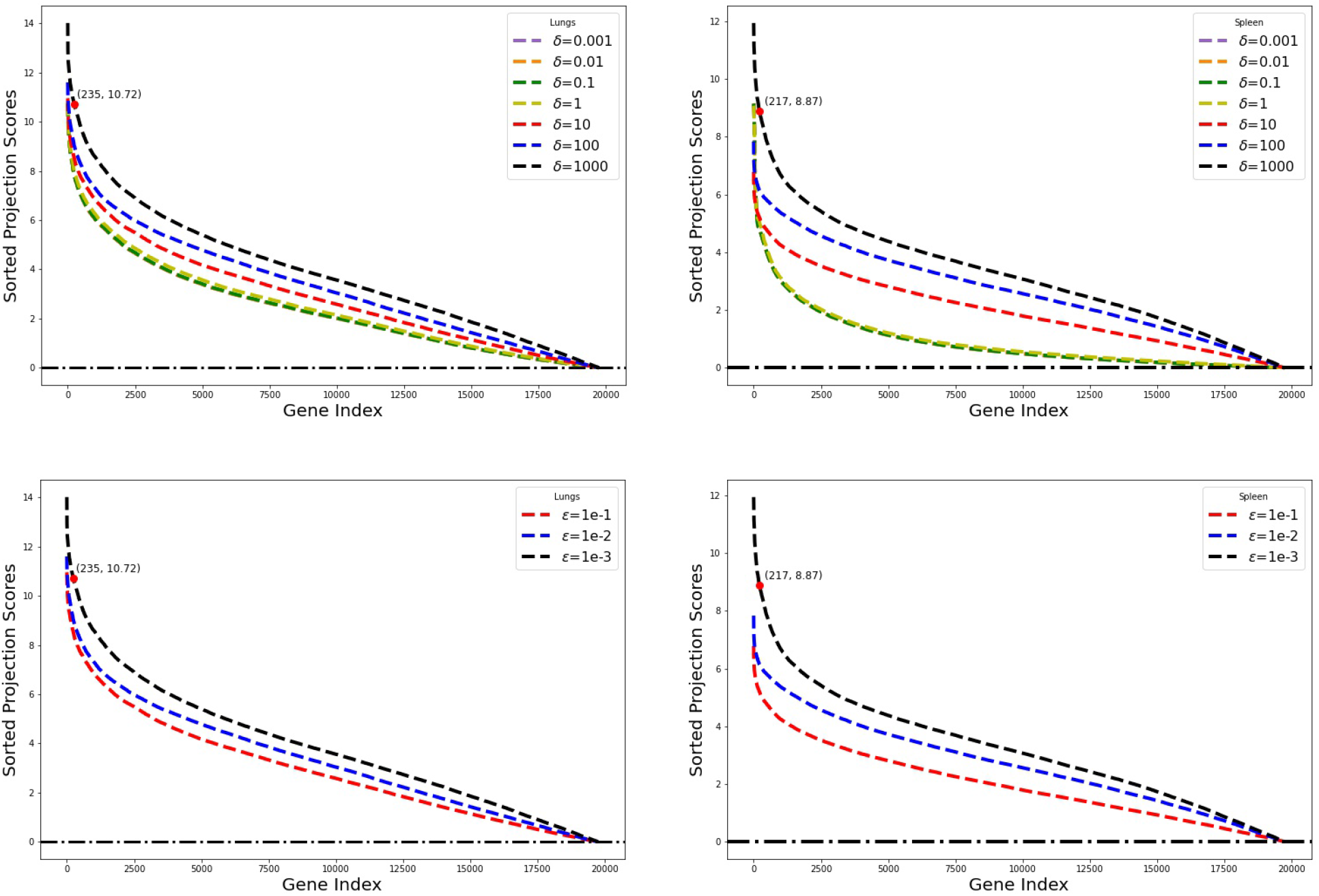
Projection scores for the lungs (L) and spleen (R) datasets corresponding to different *δ* and *ϵ* values with *M* := *D*_*ϵ*,1_(**z**)^2^.

We also compare the amount of information that the *𝓁*_1_-WGEPs capture with *M* := *D*_*ϵ*,1_(**z**)^2^ to those of *M* := *D*_*ϵ*,1_(**z**) and *M* := *I*, and found that the latter choices of *M* yield lower projection scores than the former choice for the same parameter values in Figure 1. Figures 1-3 demonstrate that regularizing (1.6) with 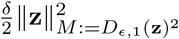 captures more informative gene expression profiles across tissues than with the custom regularization terms, 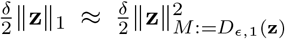 and 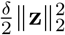. In addition, Figures 2-3 show that the projection scores corresponding to different *ϵ* and *δ* are relatively the same for *M* := *D*_*ϵ*,1_(**z**) and *M* := *D*_*ϵ*,2_(**z**), respectively.

**Figure 2:**
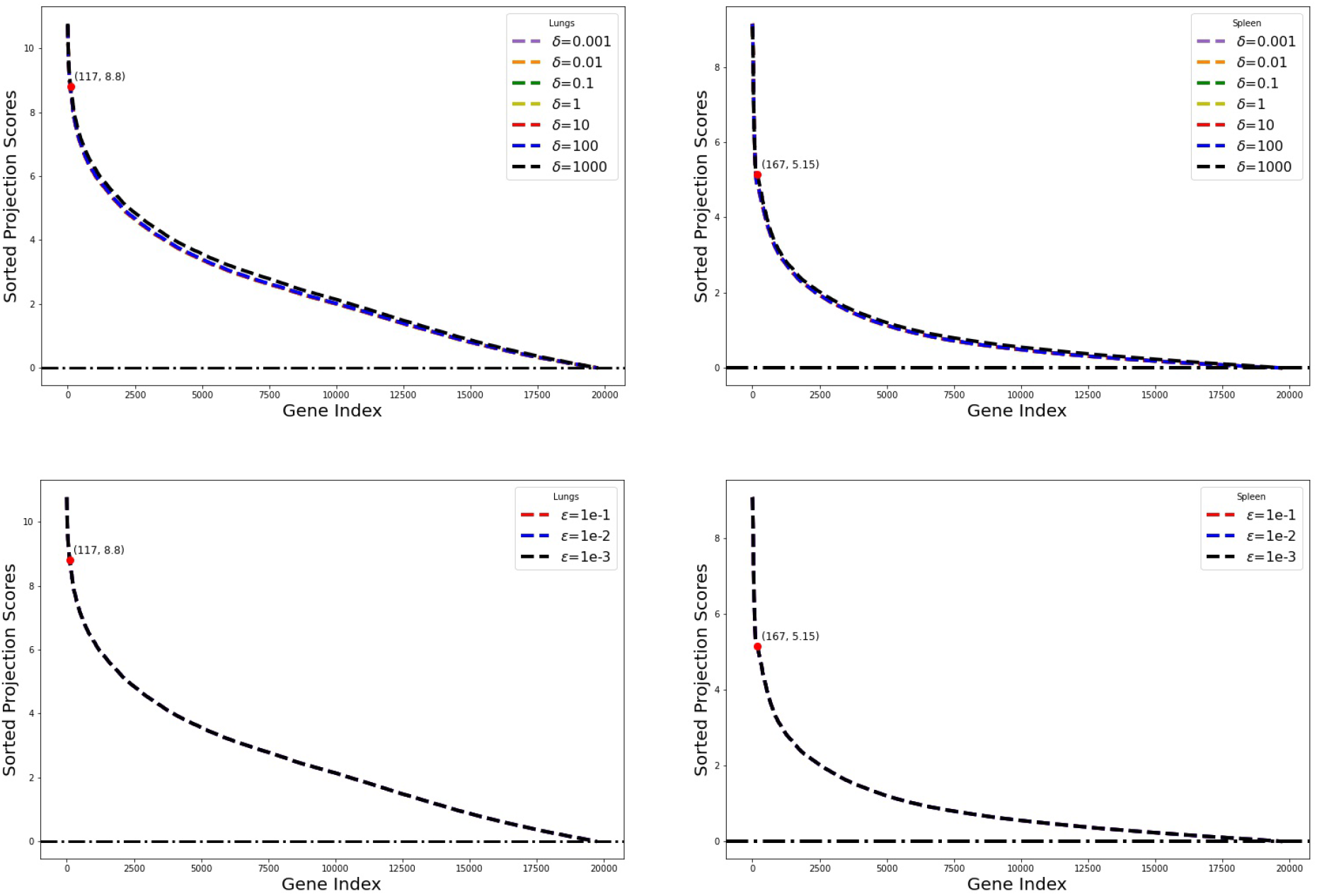
Projection scores for the lungs (L) and spleen (R) datasets corresponding to different *δ* and *ϵ* values with *M* := *D*_*ϵ*,1_(**z**).

**Figure 3:**
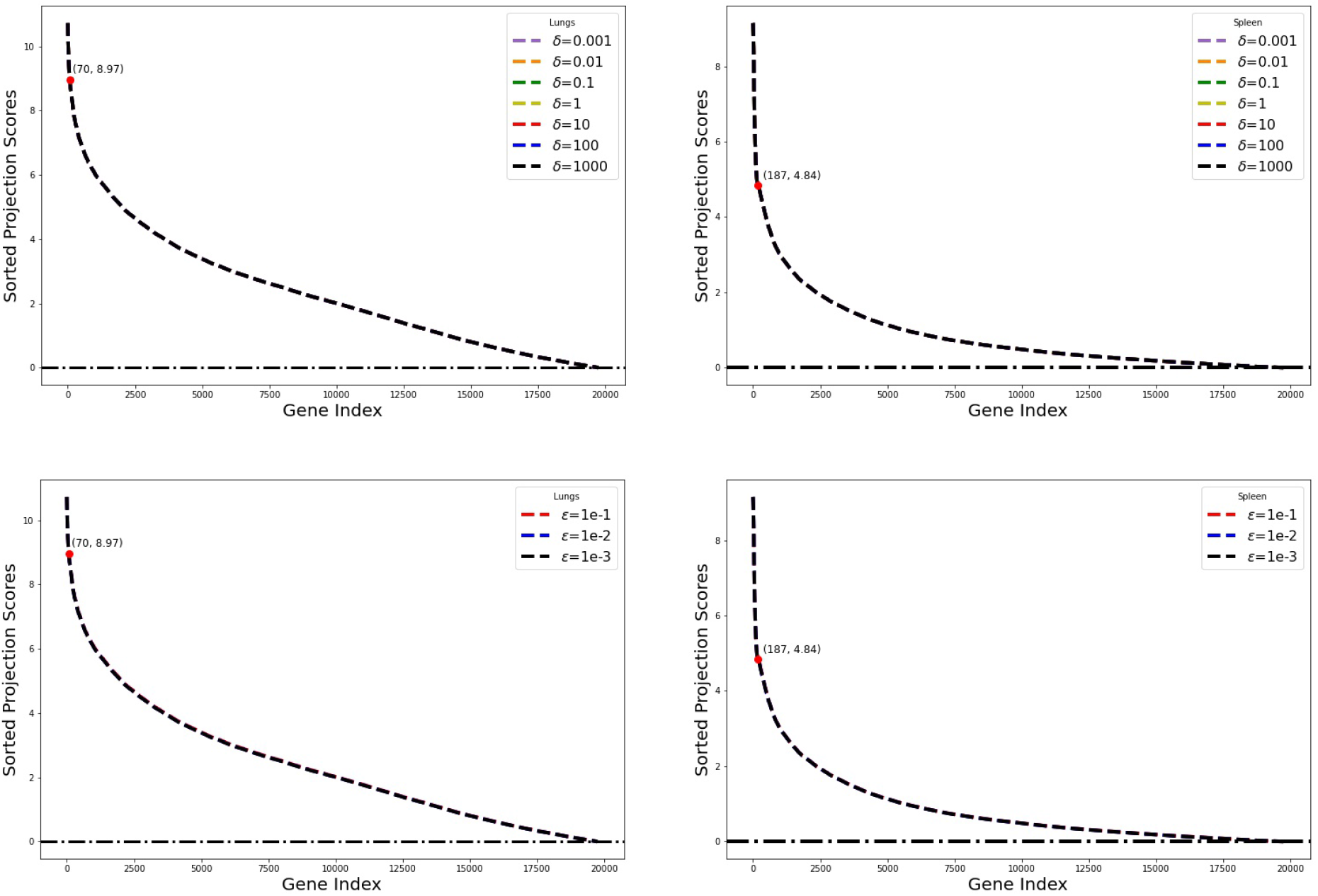
Projection scores for the lungs (L) and spleen (R) datasets corresponding to different *δ* and *ϵ* values with *M* := *D*_*ϵ*,2_(**z**).

A total of 235 and 217 genes are selected from the lungs and spleen datasets, respectively, by using the `KneeLocator` operator [33] from the `Kneed` python package. The `KneeLocator` locates the maximum curvature of the black curves in Figures 1-3. As shown in Figure 1, the maximum curvature of the ‘black’ curves occur at (235, 10.72) and (217, 8.87) for the lungs and spleen data, respectively, which are greater that those of Figures 2-3. The projection scores corresponding to the top 235 or 217 most informative genes, in both tissues, are to the left of the maximum curvatures. The gene expression profiles selected from both tissues are significantly exclusive. Figures 4 and 5 show the top 50 most informative gene expression profiles corresponding to the lungs and spleen datasets, respectively.

**Figure 4:**
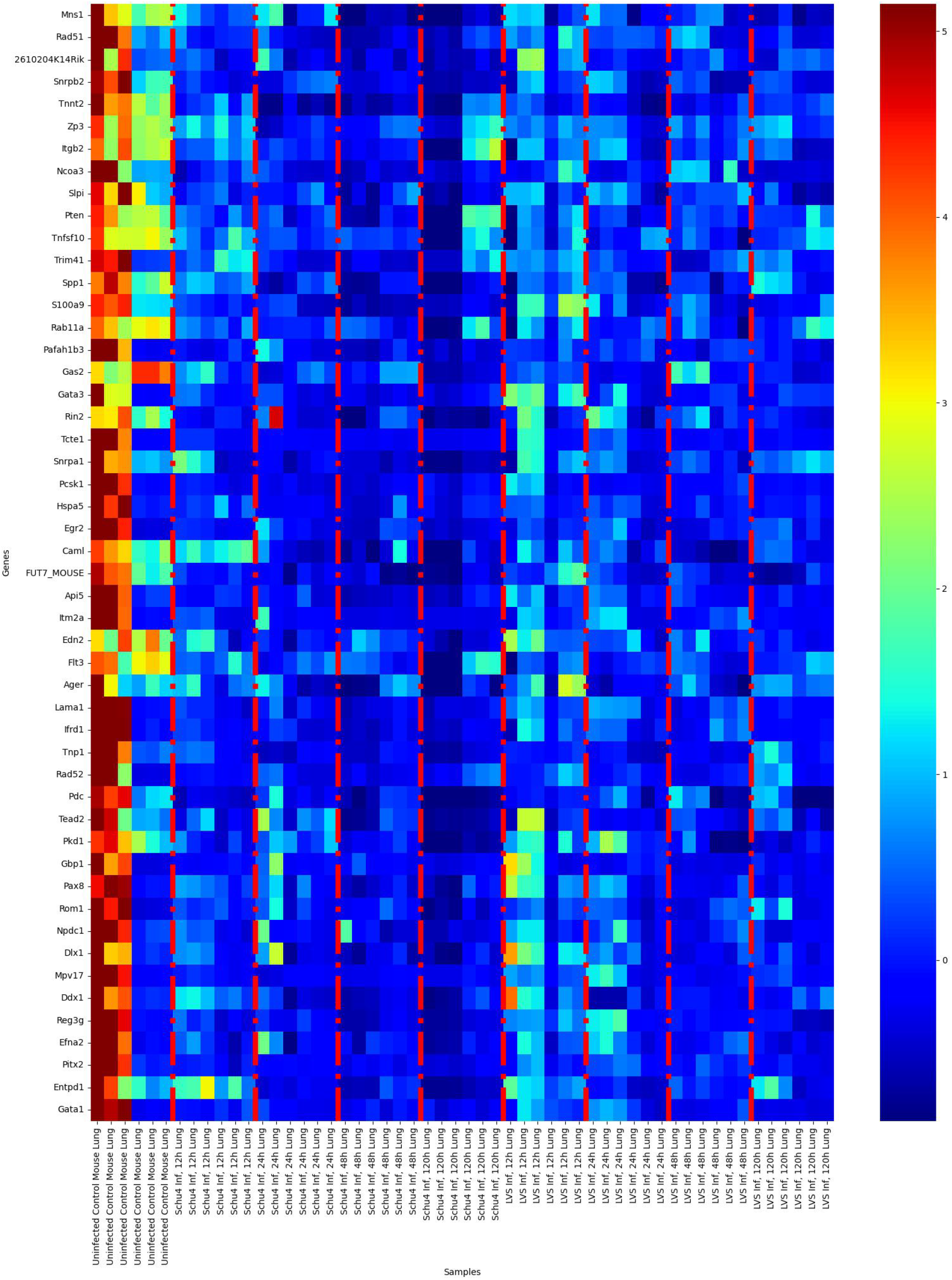
Profiles of top 50 genes that are highly significant in lungs partitioned by time points (L-R).

**Figure 5:**
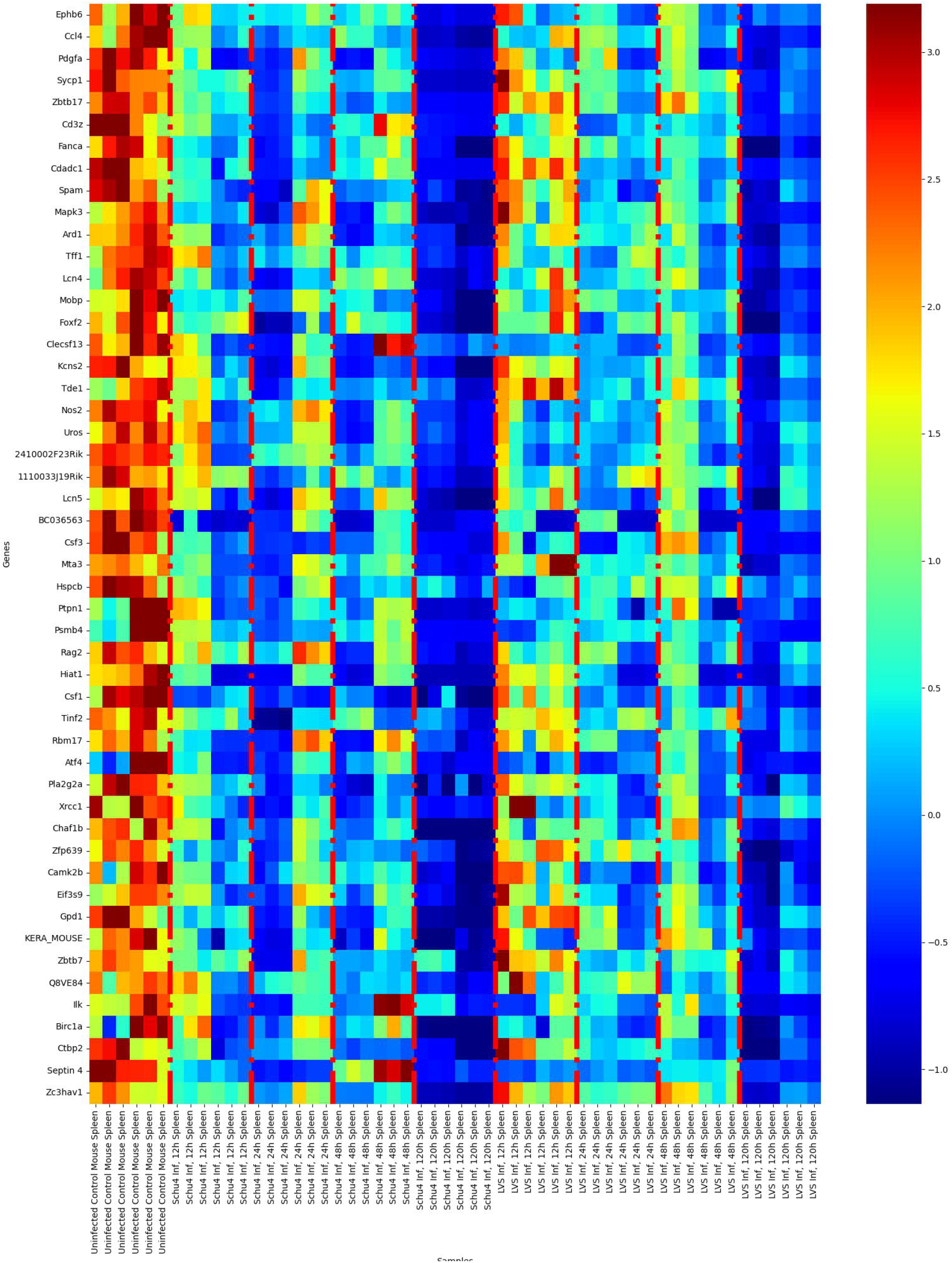
Profiles of top 50 genes that are highly significant in spleen partitioned by time points (L-R).

### 3.3 Training and Validating Transfer Learning Models with the Top 235 Most Informative Genes

In all computed examples presented in this subsection, we will use information from the lungs to model the host response patterns to bacterial dissemination to the spleen, by training ML models on the lungs host response data, and then validating them on the spleen host response data. Here, gene transcription patterns identified from the lungs are utilized to predict the gene transcription patterns in the spleen. We emphasize that our approach can be employed to overcome the challenges of limited labeled data.

Specifically, we will use the top 235 most informative genes from the lungs to train transfer learning models with uninfected controls and Schu4 or LVS samples as classes, then validate the trained models using the top 235 most significant genes from the spleen. Note that the lung transcription data with SMOTE-generated controls are used in the training process. However, SMOTE-generated controls in the spleen transcription data are not used as part of the test data but only for preprocessing and data integration purposes.

The model parameters/hyperparameters used in the training process are displayed in Table 1. For each method, the best set of parameters is determined by using `GridSearchCV` with `RepeatedStratifiedKFold(n_splits = 5, n_repeats = 10, random_state = 10`). The determined parameters are then used to train transfer learning models with the training data randomly shuffled 100 times. For the Random Forest, Logistic Regression, linear SVM, Decision Tree, KNN, AdaBoost, GradBoost, and XGBoost, the best parameters correspond to the default parameters in scikit-learn. The Naive Bayes uses 10^*−*2^ and 10^*−*3^ as variance smoothing for Schu4 and LVS datasets, respectively.

**Table 1:**
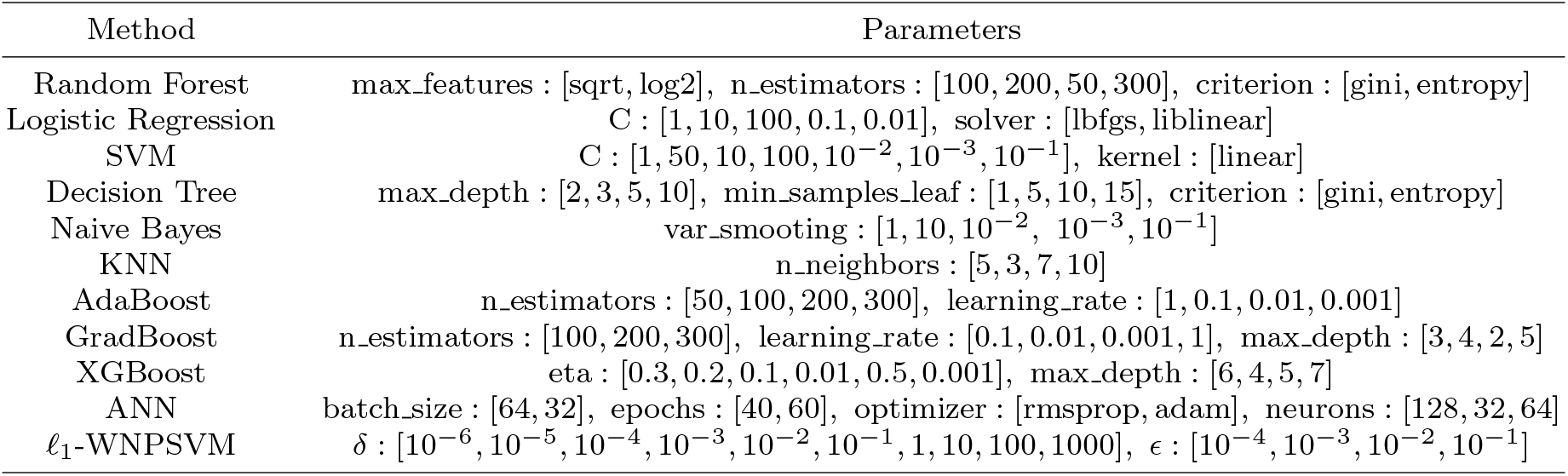
Parameters used for ML training.

For ANN, we use `TensorFlow` Keras to define a two-layer neural network with the best parameters as `batch_size = 64, epochs = 40, optimizer = “rmsprop”` and `neurons = 128`. The trained networks consist of two layers of neurons, in addition to the input layer. The hidden layer has 128 neurons while the output layer has one neuron. The activation functions of the hidden and output layers are “`ReLu`” and “`sigmoid`”, respectively.

The best parameters for the *𝓁*_1_-WNPSVM are *δ* = 10^*−*1^ and *ϵ* = 10^*−*2^ with tol = 10^*−*6^ in Algorithm 1. We will compare the performance of *𝓁*_1_-WNPSVM with the regularization matrices, *D*_*ϵ*,1_(**z**)^2^, *D*_*ϵ,q*_(**z**), *q* ∈ {0.1, 1, 2}, and *ℒ*_*i*_, *i* ∈ {1, 2}, with

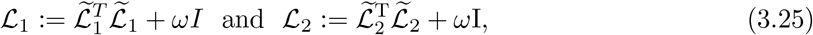

where *ω >* 0,

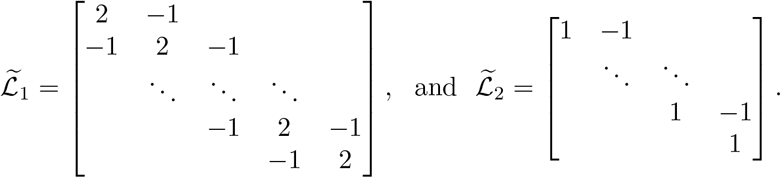

The SPD matrices *ℒ*_1_ and *ℒ*_2_ are tridiagonal matrices which are very popular in image processing applications. We will use *ω* = 3 in all computed examples. This choice of *ω* is consistent with the experiments presented in [34]. The datasets are standardized by subtracting the mean and dividing by the standard deviation.

### 3.4 Discussion of Results

Here we discuss the performance of the methods presented in Table 2. Four different examples considered below present the best and worst performing models, based on (Avg.) Bal. Acc. score. The balanced accuracy is defined in terms of the evaluation factors of the confusion matrix, i.e.,

**Table 2:**
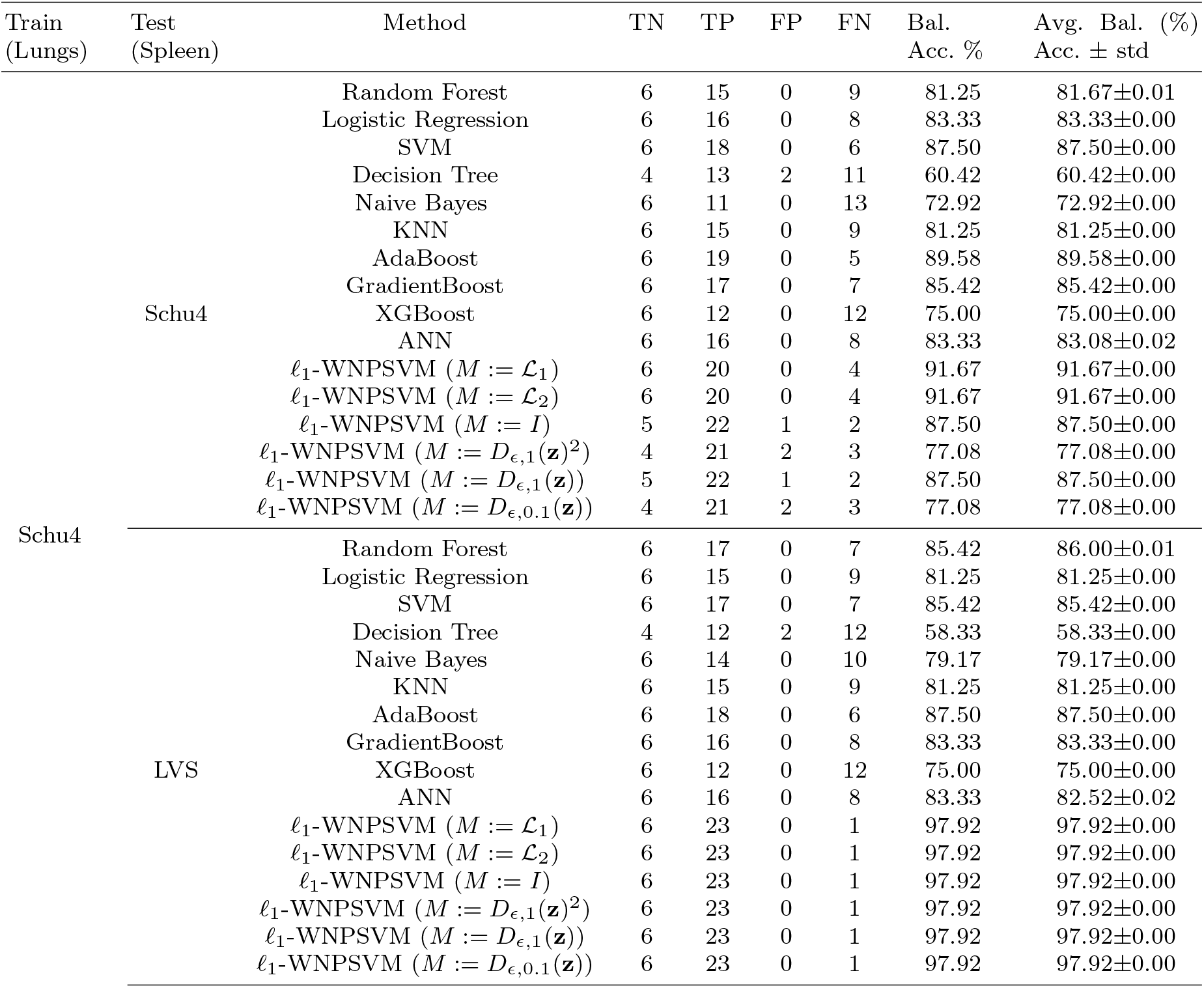
Results showing the performance of the methods with Avg. Bal. Acc. scores.

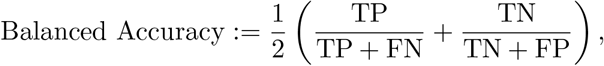

where the True Negative (TN) denotes the number of uninfected controls correctly classified, True Positive (TP) denotes the number of Schu4 or LVS samples correctly classified, False positive (FN) denotes the number of incorrectly classified Schu4 or LVS samples, and False Negative (FP) denotes the number of uninfected controls incorrectly classified. The Bal. Acc. is often considered to be well-suited for datasets with imbalanced classes.

Figure 6 shows the sorted balanced accuracy scores, for each method, over 100 different models validated on the test (spleen) data. The Average Balanced Accuracy (Avg. Bal. Acc.) scores of the methods in Figure 6 are reported in Table 2. The factors, TP, TN, FN and FP, including Bal. Acc. scores are also reported in Table 2 for transfer learning models trained with no sample shuffles. For reproducibility of the results presented therein, we use `random_state = 10`.

**Figure 6:**
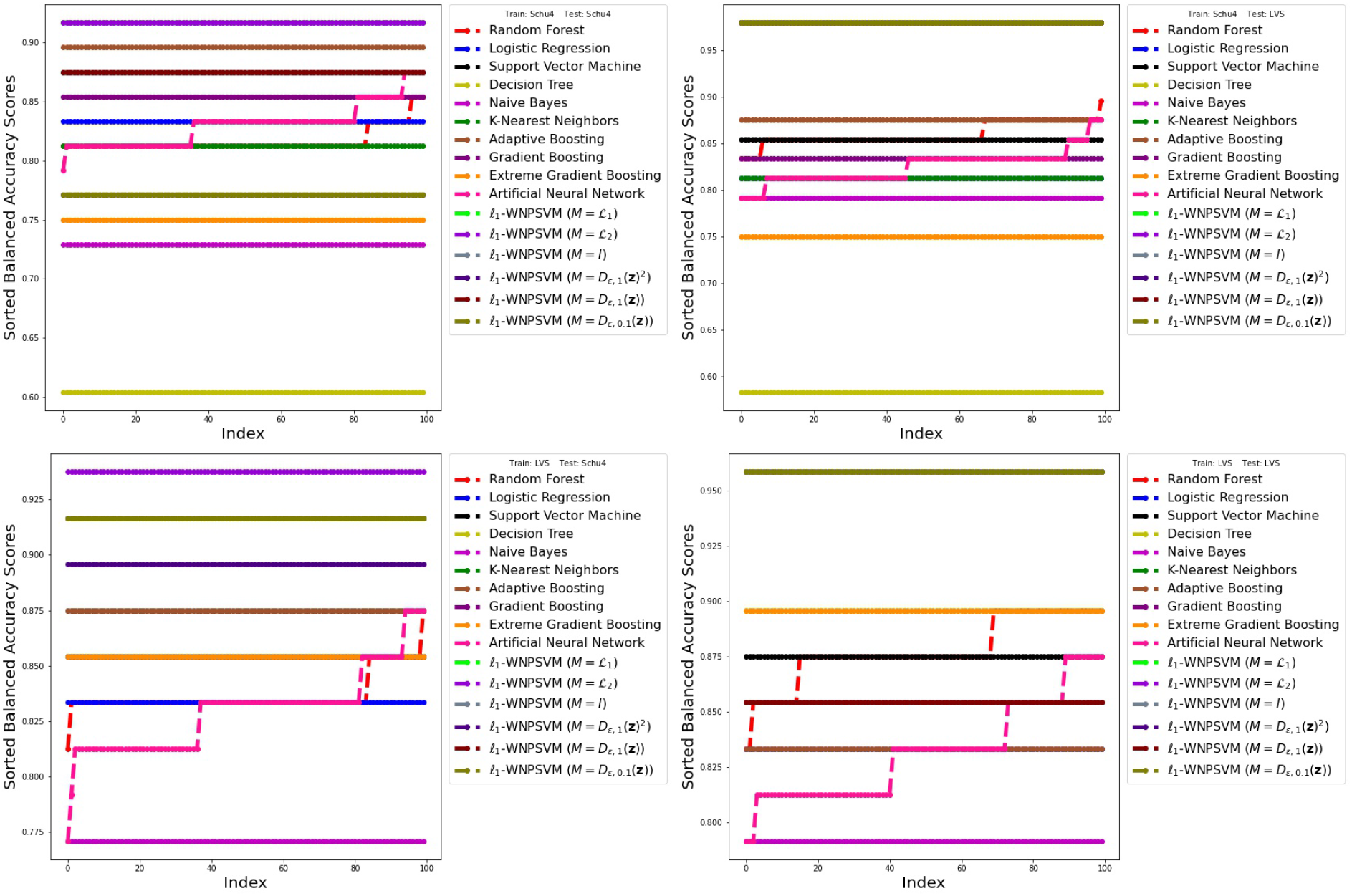
Sorted balanced accuracy scores of the methods over 100 different models validated on the test data.

**Example 3.1**. Here, transfer learning models are trained and tested on the lungs and spleen transcriptional datasets, respectively, with uninfected controls and Schu4 samples as classes. Table 2 shows that the Decision Tree is the least-performing model with an (Avg.) Bal. Acc. score of 60.42% while the *𝓁*_1_-WNPSVM with *ℒ*_*i*_, ∈ *i* {1, 2 }, in (3.25) yield the highest (Avg.) Bal. Acc. score of 91.67%.

**Example 3.2**. This example trains transfer learning models on the lungs transcriptional data with uninfected controls and Schu4 samples as classes. The trained models are tested on the uninfected controls and LVS samples from the spleen. Table 2 shows that the Decision Tree performs the least with an (Avg.) Bal. Acc. score of 58.33%, while the *𝓁*_1_-WNPSVM with the regularization matrices, *D*_*ϵ*,1_(**z**)^2^, *D*_*ϵ,q*_(**z**), *q*∈ {0.1, 1, 2 }, and *ℒ*_*i*_, *i* ∈{1, 2}, perform the best with an (Avg.) Bal. Acc. score of 97.92%.

**Example 3.3**. In this example, transfer learning models are built on the lungs transcriptional data with uninfected controls and LVS samples as classes. The trained models are applied to classify uninfected controls and Schu4 samples from the spleen. Table 3 shows that the Naive Bayes performs the worst with the least (Avg.) Bal. Acc. score of 77.08% while the *𝓁*_1_-WNPSVM with the regularization matrix *ℒ*_2_ is the best method with the highest (Avg.) Bal. Acc. score of 93.75%.

**Table 3:** Results showing the performance of the methods with Avg. Bal. Acc. scores.

| Train<br>(Lungs) | Test<br>(Spleen) | Method | TN | TP | FP | FN | Bal.<br>Acc. % | Avg. Bal. (%)<br>Acc. ± std |
| --- | --- | --- | --- | --- | --- | --- | --- | --- |
| LVS | Schu4 | Random Forest | 6 | 16 | 0 | 8 | 83.33 | 83.67±0.01 |
| | | Logistic Regression | 6 | 16 | 0 | 8 | 83.33 | 83.33±0.00 |
| | | SVM | 6 | 18 | 0 | 6 | 87.50 | 87.50±0.00 |
| | | Decision Tree | 6 | 17 | 0 | 7 | 85.42 | 85.42±0.00 |
| | | Naive Bayes | 6 | 13 | 0 | 11 | 77.08 | 77.08±0.00 |
| | | KNN | 6 | 17 | 0 | 7 | 85.42 | 85.42±0.00 |
| | | AdaBoost | 6 | 18 | 0 | 6 | 87.50 | 87.50±0.00 |
| | | GradientBoost | 6 | 17 | 0 | 7 | 85.42 | 85.42±0.00 |
| | | XGBoost | 6 | 17 | 0 | 7 | 85.42 | 85.42±0.00 |
| | | ANN | 6 | 16 | 0 | 8 | 83.33 | 83.00±0.02 |
| | | $\ell_1$-WNPSVM ($M := \mathcal{L}_1$) | 6 | 20 | 0 | 4 | 91.67 | 91.67±0.00 |
| | | $\ell_1$-WNPSVM ($M := \mathcal{L}_2$) | 6 | 21 | 0 | 3 | 93.75 | 93.75±0.00 |
| | | $\ell_1$-WNPSVM ($M := I$) | 5 | 24 | 1 | 0 | 91.67 | 91.67±0.00 |
| | | $\ell_1$-WNPSVM ($M := D_{\epsilon,1}(\mathbf{z})^2$) | 5 | 23 | 1 | 1 | 89.58 | 89.58±0.00 |
| | | $\ell_1$-WNPSVM ($M := D_{\epsilon,1}(\mathbf{z})$) | 5 | 24 | 1 | 0 | 91.67 | 91.67±0.00 |
| | | $\ell_1$-WNPSVM ($M := D_{\epsilon,0.1}(\mathbf{z})$) | 5 | 24 | 1 | 0 | 91.67 | 91.67±0.00 |
| | LVS | Random Forest | 6 | 18 | 0 | 6 | 87.50 | 87.79±0.01 |
| | | Logistic Regression | 6 | 16 | 0 | 8 | 83.33 | 83.33±0.00 |
| | | SVM | 6 | 18 | 0 | 6 | 87.50 | 87.50±0.00 |
| | | Decision Tree | 6 | 19 | 0 | 5 | 89.58 | 89.58±0.00 |
| | | Naive Bayes | 6 | 14 | 0 | 10 | 79.17 | 79.17±0.00 |
| | | KNN | 6 | 17 | 0 | 7 | 85.42 | 85.42±0.00 |
| | | AdaBoost | 6 | 16 | 0 | 8 | 83.33 | 83.33±0.00 |
| | | GradientBoost | 6 | 17 | 0 | 7 | 85.42 | 85.42±0.00 |
| | | XGBoost | 6 | 19 | 0 | 5 | 89.58 | 89.58±0.00 |
| | | ANN | 6 | 15 | 0 | 9 | 81.25 | 83.21±0.02 |
| | | $\ell_1$-WNPSVM ($M := \mathcal{L}_1$) | 6 | 22 | 0 | 2 | 95.83 | 95.83±0.00 |
| | | $\ell_1$-WNPSVM ($M := \mathcal{L}_2$) | 6 | 22 | 0 | 2 | 95.83 | 95.83±0.00 |
| | | $\ell_1$-WNPSVM ($M := I$) | 5 | 21 | 1 | 3 | 85.42 | 85.42±0.00 |
| | | $\ell_1$-WNPSVM ($M := D_{\epsilon,1}(\mathbf{z})^2$) | 6 | 22 | 0 | 2 | 95.83 | 95.83±0.00 |
| | | $\ell_1$-WNPSVM ($M := D_{\epsilon,1}(\mathbf{z})$) | 5 | 21 | 1 | 3 | 85.42 | 85.42±0.00 |
| | | $\ell_1$-WNPSVM ($M := D_{\epsilon,0.1}(\mathbf{z})$) | 6 | 22 | 0 | 2 | 95.83 | 95.83±0.00 |

**Example 3.4**. This example trains and tests transfer learning models with uninfected controls and LVS samples as classes. Table 3 shows that the Naive Bayes results in the least performing model with an (Avg.) Bal. Acc. score of 79.17% while the *𝓁*_1_-WNPSVM with the regularization matrices, *D*_*ϵ*,1_(**z**)^2^, *D*_*ϵ*,0.1_(**z**), and *ℒ*_*i*_, *i* ∈ {1, 2}, give the highest (Avg.) Bal. Acc. score of 95.83%.

Overall, the *𝓁*_1_-WNPSVM with *M* := *ℒ*_2_ is the best method in all computed examples. This illustrates the potential superiority of regularizing (1.1) with a weighted *𝓁*_2_-norm over other choices described above. We remark that the python codes for implementing the methods are available on GitHub: https://github.com/Obinnah/WNPSVM/blob/main/wnp-svm/demos/WNP_SVM.ipynb

The next section discusses pathway discovery with the selected top informative genes. Of the 235 and 217 top genes associated with the lungs and spleen datasets, respectively, only 221 and 190 gene identifiers are available for pathway analysis.

## 4 Transcriptomic Pathway Analysis

This section presents pathway analysis, also known as *functional enrichment analysis*, of available top 221 and 190 most informative gene identifiers in the lungs and spleen, respectively, by using Reactome^3^, a curated database of pathways and reactions in human biology. Our goal is to illustrate that our gene selection approach can accurately identify groups of related host genes or features that are involved in relevant biological processes such as immune response.

Reactome uses hypergeometric distribution to determine pathways that are overrepresented (enriched) in a submitted list of identifiers also referred to as *gene name list*. This overrepresentation analysis examines if a given feature list contains more genes for pathway *X* than normal by chance, then assigns a probability score, p-value, that is corrected for False Discovery Rate (FDR) using the Benjamani-Hochberg method.

In our overrepresentation analysis, all mice identifiers were converted to their human equivalent with no molecular interactors included. Reactome reports show that of the 221 most informative identifiers from the lungs data, 159 identifiers were found, and they are mapped to 210 Reactome entities (UniPort Id), with 776 pathways hitting at least one of the identifiers. As for the spleen data, 137 out of 190 identifiers were found and mapped to 189 Reactome entities while 1023 pathways were hit by at one least of the 137 identifiers. Moreover, 62 identifiers from the lungs data and 53 identifiers from the spleen data were neither found nor mapped to any entity in Reactome.

Tables 4 and 5 present the top 10 most significant pathways in the lungs and spleen, respectively, including the associated identifiers. The order of significance of the pathways is based on their p-values and FDR. Pathways with the smallest p-values and FDR, in that order, are considered to represent the most relevant features or genes.

**Table 4:** Top 10 pathways associated with the lungs data.

|  | Pathway Name | Identifiers | p-value | FDR |
| --- | --- | --- | --- | --- |
| 1. | Activation of anterior HOX genes in hindbrain development during early embryogenesis | **Egr2**, Hoxb3, Hoxd1, **Ncoa3**, Rbbp7 | $2.98 \cdot 10^{-4}$ | 0.12 |
| 2. | Activation of HOX genes during differentiation | **Egr2**, Hoxb3, Hoxd1, **Ncoa3**, Rbbp7 | $2.98 \cdot 10^{-4}$ | 0.12 |
| 3. | RUNX3 Regulates Immune Response and Cell Migration | Cbfb, **Spp1** | $8.41 \cdot 10^{-4}$ | 0.204 |
| 4. | Negative regulation of MET activity | Lig1, Ptprj, Stam | $1.00 \cdot 10^{-3}$ | 0.204 |
| 5. | TP53 Regulates Transcription of Cell Cycle Genes | Arid3a, Gadd45a, Plk2 | $1.00 \cdot 10^{-3}$ | 0.204 |
| 6. | TFAP2 (AP-2) family regulates transcription of other transcription factors | Pitx2 | $4.00 \cdot 10^{-3}$ | 0.38 |
| 7. | Regulation of gene expression in late stage (branching morphogenesis) pancreatic bud precursor cells | Hes1, Onecut1 | $6.00 \cdot 10^{-3}$ | 0.38 |
| 8. | Transcriptional regulation by RUNX3 | Brd2, Cbfb, Hes1, **Spp1**, **Tead2** | $6.00 \cdot 10^{-3}$ | 0.38 |
| 9. | STAT5 activation downstream of FLT3 ITD mutants | Flt3, Pim1 | $7.00 \cdot 10^{-3}$ | 0.38 |
| 10. | Phosphate bond hydrolysis by NUDT proteins) pancreatic bud precursor cells | Nudt1 | $7.00 \cdot 10^{-3}$ | 0.38 |

**Table 5:**
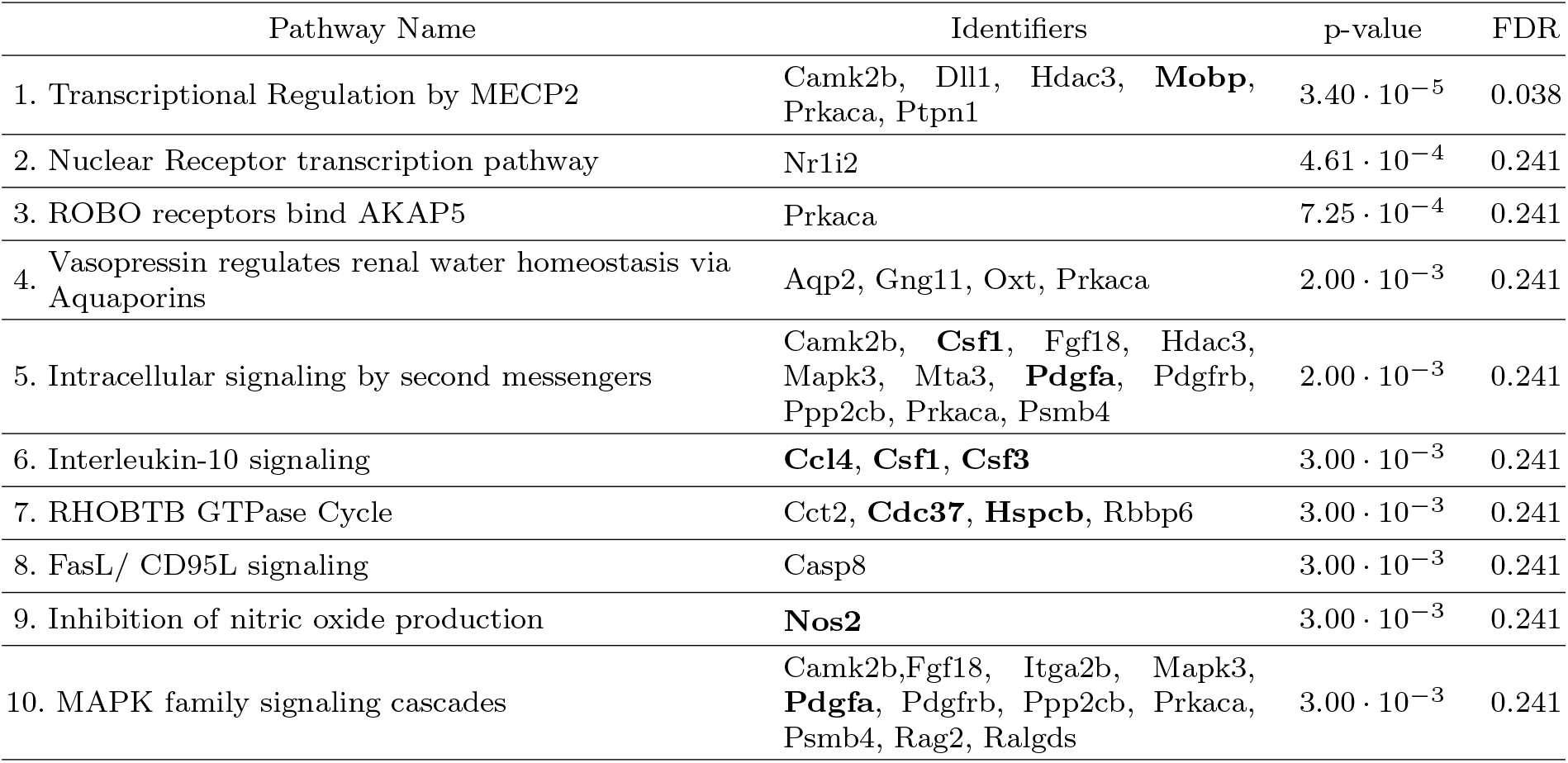
Top 10 pathways associated with the spleen data.

In Table 4, Pathways 1, 2, and 7, are categorized as developmental biology pathways while Pathways 3, 5, 6, and 8, are involved in host regulation pathways. Note that RUNX3-mediated transcription regulates development of immune system cells. Pathway 4 is a signal transduction pathway while Pathways 9 and 10, are associated with disease (cancer) and metabolism pathways, respectively. Collectively, these pathways indicate a significant host response to infection.

Similarly, in Table 5, Pathways 1 and 2, are involved in gene regulation, and Pathways 5, 7, 9, and 10, are involved in signal transduction. Pathway 6 is a central immune signaling pathway, and Pathway 8 is involved in innate immune responses to infection. Pathways 3 and 4 belong to the developmental biology pathway and transport of small molecules pathway, respectively. Identifiers highlighted in Tables 4 and 5 belong to the top 50 most informative genes in their respective datasets. Note that the expression profiles of the top 50 most informative genes are shown in Figures 4 and 5.

## 5 Conclusions

This paper aims to identify a small set of genes or features that are indicative of respiratory infection and dissemination to secondary sites. This work utilized a standardized infection model of disease and two related bacterial strains of Francisella tularensis - Schu4 and Live Vaccine Strain (LVS), with different levels of virulence that affords the opportunity to identify classifiers of infection and dissemination. Further, the use of host transcriptional data from the lungs and spleen tissues of genetically identical mice infected via the respiratory route with Schu4 and LVS allows the utilization of the identified small group of genes to perform binary classification tasks.

To achieve this objective, we propose and apply optimization and Machine Learning (ML) algorithms to analyze gene transcription datasets from the lungs and spleen tissues. We utilize weighted *𝓁*_1_-norm Generalized Eigenvalue-type Problems (*𝓁*_1_-WGEPs) to integrate the two datasets. Then dimensionally reduce their sizes by identifying the top *k* most informative host gene response profiles in both tissues associated with bacterial infection and dissemination.

The optimal solutions of *𝓁*_1_-WGEPs determine the best low-dimensional subspaces to project the datasets. The projection scores corresponding to a chosen projection direction are ranked and the utilized to select the top *k* most informative genes as biomarkers.

Consequently, the selected biomarkers from the lungs data are used to train non-parallel support vector machines incorporating transfer learning. The training process uses uninfected controls and Schu4 or LVS samples from the lungs as classes, and analogously, validate the models using the spleen data.

The performance of baseline ML algorithms, such as ANN, XGBoost, GradBoost, AdaGrad, KNN, SVM, Naive Bayes, Random Forest, Logistic Regression, and Decision Tree, are compared with our Weighted *𝓁*_1_-norm Non-Parallel Proximal Support Vector Machine (*𝓁*_1_-WNPSVM). Differently from these methods, the *𝓁*_1_-WNPSVM uses two non-parallel separating hyperplanes for binary classification. The average balanced accuracy scores of the methods over 100 folds are reported. The *𝓁*_1_-WNPSVM is the best method with a maximum (average) balanced accuracy score of 97%.

Furthermore, pathways analysis of the most significant genes in lung and spleen tissues are examined for relevant biological pathways that may aid in not only biomarkers of infection, dissemination and disease progression, but also provides information about the host response to infection that can inform the development of host-directed therapeutics. Our analysis can lead to a better understanding of infectious disease progression as well as the development of biomarkers that are able to predict clinical outcomes of disease during treatment.

## Footnotes

1 https://emergency.cdc.gov/agent/tularemia/faq.asp

2 https://www.mathworks.com/matlabcentral/fileexchange/75401-synthetic-minority-over-sampling-tech\nique-smote

3 https://reactome.org/

## Notes

### Competing Interest Statement

The authors have declared no competing interest.

